# Neural activity in the anterior cingulate cortex is required for effort-based decision making

**DOI:** 10.1101/2022.03.22.485350

**Authors:** Adrienne Q. Kashay, Jovian Y. Cheung, Rahil N. Vaknalli, Molly J. Delaney, Michael B. Navarro, Christabelle Junaidi, Faith Veenker, Morgan E. Neuwirth, Christopher J. Gabriel, Laura A. DeNardo, Scott A. Wilke

## Abstract

Adaptive decision making requires the evaluation of cost-benefit tradeoffs to guide action selection. Effort-based decision making (EBD) involves weighing predicted gains against effort costs and is disrupted in several neuropsychiatric disorders. The anterior cingulate cortex (ACC) is postulated to control effort-based choice via its role in encoding the value of overcoming effort costs in rodent EBD assays. However, temporally precise methods of manipulating neural activity have rarely been applied to EBD. We developed and validated a mouse version of the barrier T-maze EBD task, in which action selection is spatiotemporally segregated from surmounting an effortful obstacle and effort is minimally confounded with time cost. Optogenetic silencing of ACC excitatory neurons during action selection, rapidly and reversibly impaired preference to exert greater effort for a larger reward, when a less effortful alternative was available. Detailed analysis of mouse choice trajectories revealed that silencing ACC disrupts kinematics, especially prior to high effort choices. However, there were no effects on overall mobility or tendency to exert effort in non-choice assays. These findings establish causality between ACC neural activity and effortful action selection during spatial cost-benefit decision making.

**SIGNIFICANCE STATEMENT:** Disturbances in evaluating effort-based costs during decision making occur in depression, schizophrenia, addiction and Parkinson’s disease. Precisely resolving the function of prefrontal brain regions in mediating these processes will reveal key loci of dysfunction and potential therapeutic intervention in these disorders.

## INTRODUCTION

Goal-directed decision making requires prospectively evaluating cost-benefit tradeoffs to guide action selection. Effort costs, the motor and cognitive resources required for performing an action, are a critical consideration in making adaptive choices. Effort-based decision making (EBD) is specifically impaired in patients with depression, schizophrenia, addiction and Parkinson’s disease^1–6^. The anterior cingulate cortex (ACC) is a prefrontal region that has consistently been implicated in evaluating both cognitive and physical effort costs^7–15^. Moreover, the ACC has been associated with a wide variety of behavioral functions relevant to decision making more broadly. These include computation of reward value and action outcomes, representing reward prediction errors, model-based action selection, exploration of alternative strategies, tracking outcomes across trials, and others^16–25^. Thus, the ACC is well positioned to mediate effort-based decision making including evaluating the effort-related value of potential actions, selecting between these options and carrying them to completion.

Much of the seminal work on the neurobiology of spatial EBD has been done using the barrier T-maze task, originally developed by Salamone and colleagues^26,27^. In this task, animals must choose between climbing a vertical barrier to access a large reward or taking an unobstructed path to a smaller reward. Lesion studies, done primarily in rats, have found that ACC, but not prelimbic or orbitofrontal regions, disrupts spatial EBD^7,8,13,28,29^ (though see a null result in mice^30^). However, ACC lesions or inactivation do not universally impair effortful behavior or effortful choice tasks^13,29^. For example, operant lever-pressing choice tasks have yielded both negative results^13^ and findings suggesting a reduced preference to exert effort for larger rewards^31–33^. This work clearly implicates ACC in rodent EBD, but studies have primarily used lesions or pharmacology to establish causal linkages. These approaches generally affect larger regions, produce observations lacking temporal precision and cannot address cell-type specific contributions in ACC.

The temporal relationship between the activity of ACC neurons and effort-based decision making has been explored using electrophysiological approaches. Recordings of population or single neuron activity from ACC find representations of choice options with the highest utility based on integrating effort cost and reward value^11,14,34,35^. These studies find clear encoding of effort-related value, but also high degrees of heterogeneity indicating that ACC neurons can respond to a broad set of task features. Other studies have begun to explore the encoding of effort-related behaviors at the ensemble level using miniaturized head-mounted microscopes, or multielectrode recordings^32,36^. Similarly, local field potential recordings between ACC and reward related regions implicate oscillatory activity in spatial effort-based decision making^37–39^. Thus, ACC is critical for representing the effort-related value of actions and activity is temporally aligned with specific aspects of decision making.

Here, we overcome some of the limitations of traditional lesion and pharmacologic methods used to study the causal role of ACC in EBD. Because prior studies used methods lacking temporal precision, the question of when during a trial ACC is required for EBD is not known. Moreover, when ACC function is disrupted, it is not known how this impacts the action sequences required to select and carry out a choice. In this study we resolve these questions in order to shed light on the mechanistic function of ACC in EBD. We use mice to clarify conflicting results as to the role of ACC for EBD across species and to enable work in a genetically tractable model. To do this, we developed a protocol for training and testing mice in the barrier T-maze. Mice readily learn the task and behavior is sensitive to altering choice utility by changing reward magnitude or effort cost with minimal confounding due to time cost. We then show that bilateral optogenetic silencing of ACC excitatory neurons prior to or during action selection can rapidly and reversibly disrupt preference for high effort, high reward choices on a trial-by-trial basis. When effort cost was equalized, mice switched to choosing the more rewarded arm on all trials, indicating that spatial memory, reward preference and ability to enact effortful behavior were not disrupted. We then used machine learning based behavior tracking and custom zone-based analysis software to investigate how silencing ACC affects choice trajectory during EBD. Silencing ACC disrupts the microstructure of choice behavior, introducing brief pauses as mice approach the maze choice point suggesting that ACC mediates online control over ballistic movements during EBD. These effects were specific to an explicitly goal-directed context as silencing ACC did not disrupt effort or movement when overt decision making is not required.

## MATERIALS AND METHODS

All experiments were conducted in accordance with procedures established by the administrative panels on laboratory animal care at the University of California, Los Angeles.

### Animals

All experiments use C57BL/6J wildtype mice of either sex (12-18 weeks of age), bred in house or ordered from Jackson Laboratories.

### Viral vectors and targeting/expression

Adult mice were stereotaxically injected at 7-10 weeks of age with 600 nl of 1.5×10^13^ vg/ml AAV1-CKIIa-stGtACR2-FusionRed (AddGene) or 7×10^12^ vg/ml AAV1-CKIIa-mCherry control at bilateral ACC (antero-posterior (AP) +1.4, medio-lateral (ML) ±0.35, dorso-ventral (DV) −1.3mm relative to bregma). Bilateral 200 μm diameter, 0.37NA fiber optic cannulae (Newdoon) were implanted over ACC at +1.4 AP, ±0.35 ML, −1.3 DV. Implants were affixed onto the skull using Metabond dental cement (Parkell). Targeting of viral constructs/cannulae were confirmed via post-hoc histology without immunostaining.

### T-maze Behavior Apparatus

Our T-Maze was constructed in-house using white corrugated plastic (Coroplast). Removable, transparent food cups were inserted into slots in each arm. Barriers were constructed from diamond pattern wire-mesh and were 5 or 10 cm in vertical height with ~45 degrees sloping ramp on the backside and placed at a standard position in the maze. Removable guillotine style doors were inserted to create a start box, and to block off one arm of the maze during forced trials or after mice made a choice on a given trial. Experiments were conducted inside specialized chambers with an open front, LED lighting and ports to pass cabling in and out. The maze was fixed in place within the chamber and a webcam was mounted overhead in a fixed location (ELP 2.8-12mm Varifocal 1.3 megapixels, 30 fps). Mazes were hand-operated by an experimenter standing in front of the chamber.

### T-maze Behavioral Procedures

Mice were housed under standard 12-h light/dark cycles, and all experiments were performed during the light portion of the cycle. Mice were group housed and allowed to fully recover post-operatively (~1 week). Mice were habituated to human handling and food intake was restricted until mice were 80-85% of the *ad libitum* feeding weight. Throughout the experiment, mice were weighed and fed daily rations. Reward pellets were peanut butter chips (Reese’s), precisely cut into small cubes (~0.01g; tolerance 0.08-0.12g). A random subset of pellets were weighed to ensure consistency across batches. After mice reached target weight, they are allowed to explore the maze with 20 food pellets scattered throughout. Once mice ate all the food pellets in <5 minutes, behavior was further shaped so that mice ran trials and could easily locate single food pellets in each reward cup. Thereafter, one maze arm was designated as high reward (HR, 3 pellets) and the other as low reward (LR, 1 pellet). The HR arm was counterbalanced (left vs. right) across mice, with mice trained against any inherent initial side biases. On each training day, mice were forced to sample each arm and then ran 15 choice trials. Once mice learned to choose the HR arm (>70%, ~2-7days), barrier training commenced. Mice were trained on a 5cm, then 10cm barrier in the HR arm. If mice refused to climb the barrier, additional forced trials were introduced to ensure adequate sampling of HR arm (up to 4/day). Training was complete when mice achieved >70% high effort/HR choices on 2 consecutive days (~5-10 days). For some experiments, the reward differential was manipulated and/or a second 10cm barrier was introduced in the LR arm to equalize effort cost between choices. Mice ran behavior trials 5 days per week, with food deprivation maintained during the weekend. After each trial, mice were placed in a home cage during the inter-trial interval (ITI), while the maze was reset (timed to ~60 seconds). Trials were run consecutively for each individual mouse and the maze was cleaned thoroughly with 70% ethanol in between mice. For optogenetics experiments we utilized white barriers and transparent food cups to enhance contrast between subject and background to facilitate segmentation of animals during behavior.

### Tail suspension test (TST)

For the TST, mice were suspended by taping their tail to a bar ~12 inches above the tabletop, within 4 inch wide stalls of a custom apparatus constructed for this purpose. A 1.5 ml microcentrifuge tube was cut and placed over the tail base to prevent the mouse from climbing its tail during testing. Per standard protocols, and to ensure a stable base level of struggling, the first 2 minutes of the TST was not analyzed. The TST for optogenetics experiments was 14 minutes in duration and only one mouse was run at a time. Laser light for optogenetics was delivered in 4x, 3 minute blocks (OFF-ON-OFF-ON) with laser stimulation parameters as per below. The apparatus was thoroughly cleaned between uses. TST videos were analyzed manually on a per minute basis by an experimenter that was blinded to the conditions being tested and using an established method^40^.

### Open field (OF)

OF testing was conducted in a standard sized arena (40×40cm), with a height of 30cm on each side. Video was collected from above and subsequent analysis was done using Bioviewer software to automatically track mice in the maze. Open field testing was 8 minutes in duration and the mouse was placed by hand into the lower right corner of the apparatus. Optogenetic stimulation was delivered in 4x, 2 minute blocks (OFF-ON-OFF-ON) with laser stimulation parameters as per below. Mean velocity in the apparatus, during total light OFF vs. ON epochs was analyzed as a measure of overall activity level. The OF apparatus was cleaned thoroughly between uses.

### Optogenetic Protocols

For optogenetic silencing during behavior, light stimulation was delivered via a fiberoptic cable fed through a rotary joint commutator to a bifurcated optogenetic patch cable (Doric Lenses), and attached to a 150 mW, 473 nm DPSS fiber coupled laser (Shanghai Lasercentury). For optogenetic silencing the laser was triggered with commercial software and an external pulse generator (Prizmatix), or triggered manually. Light power was measured at the beginning of each behavior session using a light power meter (Thorlabs), measured as 5mW total light power (2.5 mW/side). Light OFF and ON trials were interspersed pseudorandomly within each set of behavior trials with light delivered with spatiotemporal precision relative to T-maze zones. For some experiments the laser was turned on and off manually based on the location of the mouse in the maze or at specific times during the intertrial-interval (ITI) or in the start box. Mice were maintained in the start box of the maze for 6 seconds before opening the door to initiate a trial. For all optogenetic experiments, mice first ran two forced trials to either arm, followed by 16 choice trials each day (8 light OFF and 8 light ON). We delivered blue laser light to optogenetically silence the bilateral ACC at specific times during the ITI, while the mouse was in the start box of the maze, or during a period extending from start gate through the choice point of the T-maze. Light was delivered during discrete periods designated for optogenetic testing. Mice were run up to 3 consecutive days with a given maze arrangement. There was no apparent change in responsiveness to silencing over the course of the experiments. T-maze experiments ended with 1-3 days in which a second barrier was inserted to equalize effort between conditions. Because control mice typically chose the HR barrier arm on most trials, we did not run them on the 2-barrier condition. The OF and TST were performed as described above to examine effects of optogenetic silencing on overall effort and mobility.

### Behavior Analyses

In all cases, the experimenter manually recorded the choice of mice in the T-maze, and these data were cross-checked against recorded videos to verify accuracy, and correct timing of laser triggering. These choice data are reflected in plots of %HR choice throughout. In our initial validation dataset, we manually annotated videos to determine the length of each choice trial from the time point just after the start until the point at which the tip of the mouse’s nose reached the reward cup in either arm. For subsequent optogenetic experiments, we employed a more detailed, automated analysis of choice trials. This analysis was not done for the validation data due to difficulties with automated tracking of mice in the original version of the maze. For automated analyses we first used DeepLabCut (DLC)^41^ to train a neural network to track multiple points on the mouse during choice trials. We used a point centered on the implant as our main tracking point for subsequent analysis of time in zones for mice in distinct parts of the maze. Maze zones were defined in a consistent manner based on standardized markings on the maze itself. The barrier, reward cups and start gate were placed in identical locations for each trial based on specialized slots or marks on the outside of the maze. We used BehaviorDEPOT^42^ software and custom MATLAB code to define the choice made on each trial and to divide the maze into zones for further analysis. We calculated time in each zone and parsed these based on choice (HR vs. LR) and laser status (light OFF vs. ON). The microstructure of choice trajectories was analyzed manually from behavior videos on a frame-by-frame basis with a pause defined as >2 frames in which mice did not progress towards goal.

### Statistics

Unless otherwise specified, nonparametric tests or ANOVA was used to assess significance. Statistics were calculated using custom MATLAB code or Graphpad Prism. All statistical parameters, including test statistics, correction for multiple comparisons, and sampling of repeated measurements are stated in the main figure legends. Error bars represent ±SEM. The code and data used to generate and support the findings of this study are available from the corresponding author upon reasonable request.

## RESULTS

### Behavioral Validation

Most studies exploring the role of the ACC in spatial, effort-based choice tasks used rats. To investigate the role of the ACC in mouse models, we designed and validated a barrier T-maze training and testing procedure for use in mice (see methods). We built a custom, scaled down version of a standard rat T-maze, with manually removable guillotine style doors, wire mesh barriers of different sizes and transparent food cups to hold reward pellets (**Fig. 1A and B**). Mice were food deprived, habituated to human handling and behavior was shaped such that mice sought food in the reward wells. Next, animals were trained using 3 reward pellets in a high reward (HR) arm and one pellet in the low reward (LR) arm (**Fig. 1A**). Mice were trained to a threshold of >70% HR choices, first without a barrier and then sequentially with 5 and 10 cm wire mesh barriers before initiating experimental testing procedures (**Fig. 1C**). Mice initially ran 1 forced trial to each arm, followed by 16 choice trials during training and testing.

**Figure 1.**
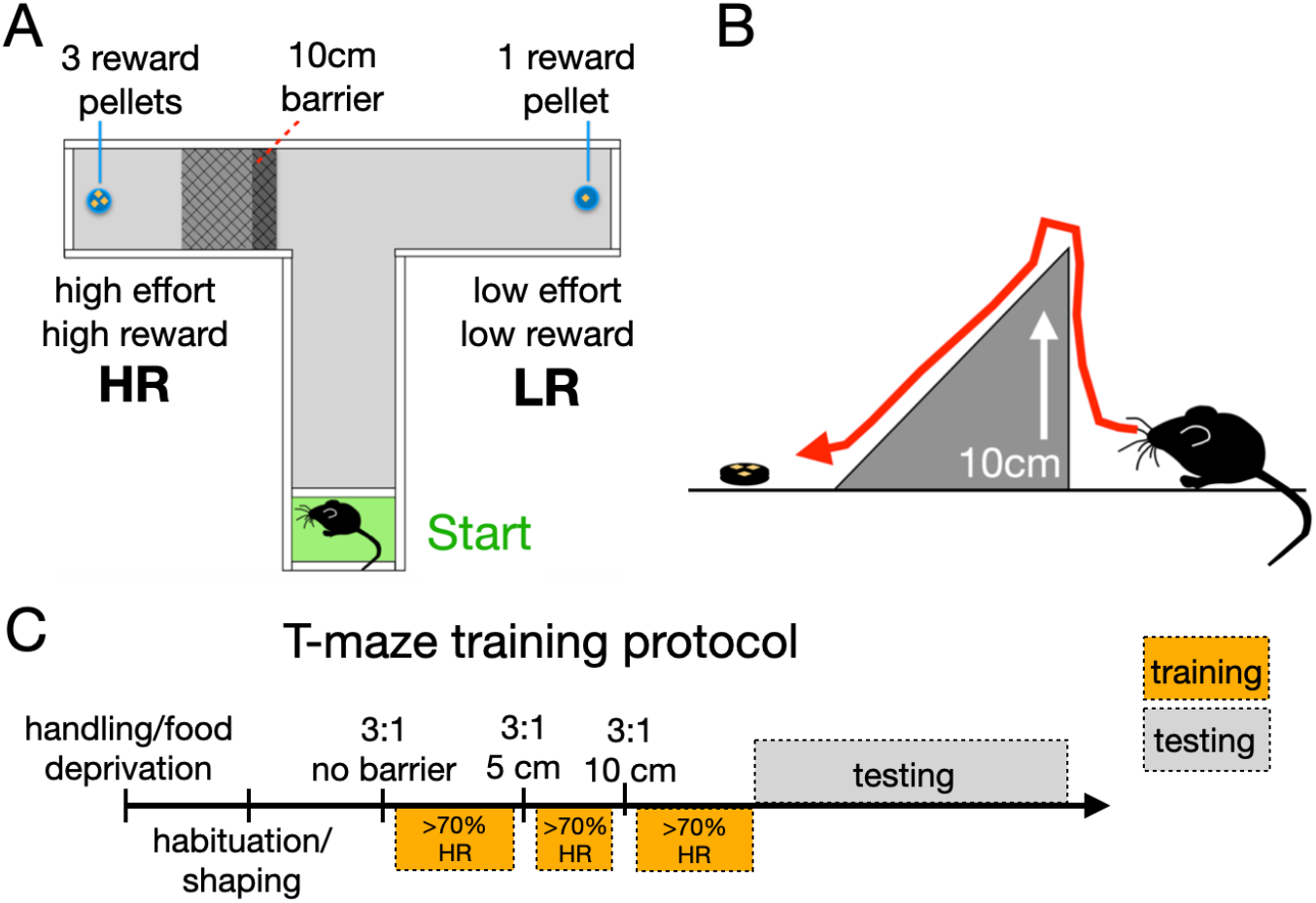
Mouse barrier T-maze schematic and training protocol. **A**) Barrier T-maze design with 10cm wire mesh barrier, and 3:1 reward differential for high effort, high reward (HR) vs. low effort, low reward (LR) arms. **B**) Side view of barrier showing 10 cm vertical side with 45 degree slope down to reward cup. **C**) Overview of general approach for training and testing mice in the assay - Following habituation/shaping, mice are trained to >70% HR choices with no barrier, 5cm barrier and 10cm barriers preceding testing phase.

To validate that the behavior of mice in our T-maze assay was sensitive to manipulations of reward differential between HR vs. LR arms or effort cost (barrier), we performed an initial validation experiment in wildtype mice (**Fig. 2**). Following training, we systematically changed the number of HR reward pellets and/or the effort associated with HR vs. LR arms of the maze and recorded the resulting pattern of choices (**Fig. 2A and B**).

**Figure 2.**
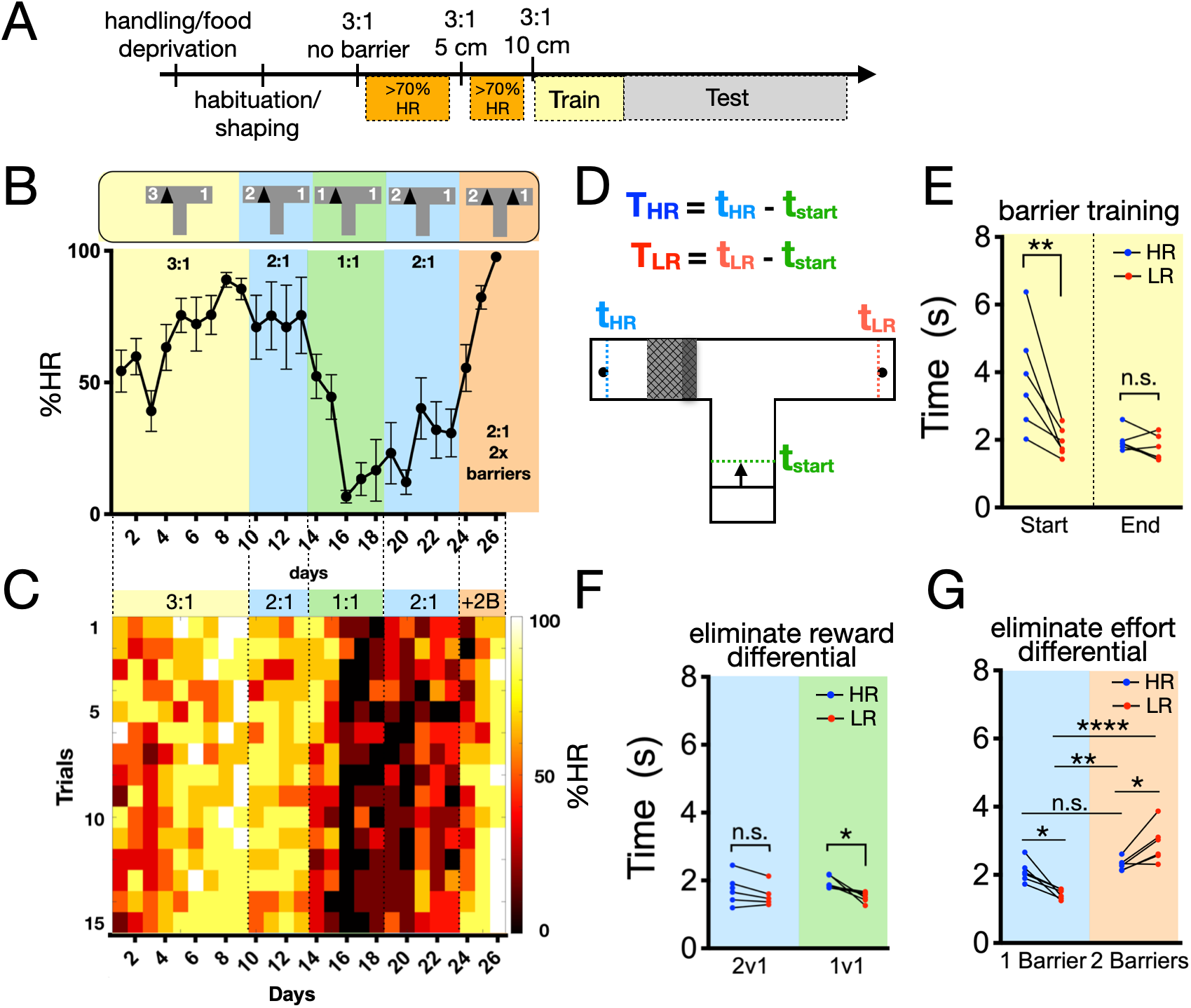
Validation of T-maze training and testing protocol in mice. **A**) For validation experiment, resutls are shown for the 3:1 +10cm barrier training portion of the task, followed by manipulations of reward differential or barrier arrangement (Test). **B**) Outcome of training and testing phase shown as %HR choices. Reward in high effort arm is systematically changed, and a second barrier is added to equalize effort in the final phase. **C**) Same data as B, but shown with mean %HR on a per trial basis to reveal within session changes. To examine behavior in greater detail, we calculated the time it takes mice to complete HR and LR choices (from start of a trial through reaching reward cup). **D**) Schematic representation of how trial lengths were calculated for **E-G. E**) Quantification of trial length for start vs. end of barrier training, [*F_Interaction_*(1,20)=5.84, *P*=0.025, *F_Time_*(1,20)=9.21, *P*=0.007, *F_Choice_*(1,20)=8.58, *P*=0.008], 2-way ANOVA with Sidak posthoc test. **F**) Quantification of trial length before vs. after eliminating reward differential (2v1-to-1v1), [*F_Interaction_*(1,19)=1.20, *P*=0.288, *F_Reward_*(1,19)=0.363, *P*=0.55, *F_Choice_*(1,19)=5.98, *P*=0.024], 2-way ANOVA with Sidak posthoc test. **G**) Quantification of trial length before vs. after eliminating effort differential (1Barrier-to-2Barriers), [*F_Interaction_*(1,20)=20.97, *P*=0.0002, *F_Barrier_*(1,20)=37.14, *P*<0.0001, *F_Choice_*(1,20)=0.037, *P*=0.85], 2-way ANOVA with Sidak posthoc test. N=6 mice, n.s. = not significant, *p<0.05, **p<0.01, ***p<.001, ****p<0.0001.

For consistency, we have used ‘HR’ to indicate the original high effort, high reward arm of the maze throughout (**Fig. 2B**). Mice initially chose the high effort, high reward arm ~50% of the time, gradually approaching ~80% HR choices (**Fig. 2B**). Reducing the reward differential (from 3:1 to 2:1 to 1:1) resulted in mice making fewer HR choices, particularly after changing to a 1:1 reward ratio (no differential) such that there was no additional utility associated with climbing the barrier (**Fig. 2B**). HR choices increased slightly after reinstating a 2:1 reward differential, though remained lower than during the initial 2:1 stage (**Fig. 2B**). These data indicate that mice adjust their willingness to exert effort in proportion to the magnitude of associated reward (**Fig. 2B**). The lower proportion of HR choices after reinstating a reward differential suggests that choices are partially influenced by recent history of reward across recent days. Finally, in order to determine whether mice are sensitive to changes in the amount of effort, independent of reward differential, we inserted a second 10 cm barrier in the LR arm of the maze. When effort cost was equivalent, mice rapidly adjusted their strategy to choose the more highly rewarded arm on nearly every trial (**Fig. 2B**). Thus, mice in our assay actively organize their behavior to optimize the utility of choices based on effort-reward tradeoffs. Learning of new contingencies could be observed within-session, but also appear to consolidate across days (**Fig. 2C**).

The time required for decision making is a relevant variable that may reflect differential engagement of distinct systems for action selection^43,44^. We hypothesized that if mice are making deliberative, goal-directed choices then this would be reflected in the timing of choice trials. Thus, we calculated trial length using overhead videos collected during EBD (**Fig. 2D**). At the start of 10 cm barrier training mice were significantly slower to complete high effort, HR trials compared to LR trials (**Fig. 2E**). However, at the end of training when mice approach asymptotic performance there was no difference in time required to complete trials regardless of choice type (**Fig. 2E**). Importantly, this indicates that the additional time cost incurred by climbing the barrier is minimal once mice are well trained on the task. This underscore the value of this assay for disambiguating effort from time-related costs which is a known issue with operant effort-based decision making tasks that require lever pressing^45^. Subsequent changes in reward differential had minimal impact on trial length, though a slight difference between HR and LR trial lengths did emerge when contingencies had recently changed (**Fig. 2F and G**). This may reflect an increase in cognitive effort or deliberative processing required to overcome an effort cost when outcomes are dynamic or uncertain. Finally, making effort equivalent by adding a second barrier resulted in a significant increase in LR choice time (**Fig. 2G**). Overall, these data demonstrate that mice pursue goal-directed choice strategies in our assay that are highly sensitive to relative reward magnitude and effort cost. Thus, this approach is a valid strategy for studying the neural basis of EBD in mouse models.

### Optogenetic silencing of the ACC disrupts effort-based decision making

To investigate the spatiotemporal requirement for ACC activity during EBD we expressed an inhibitory opsin (AAV-CKII-stGtACR2) in bilateral ACC and silenced excitatory neurons by delivering blue light via implanted cannulae (**Fig. 3A-C**). This approach consistently targeted a relatively restricted region of dorsal, anterior ACC (Cg1; **Fig. 3E**). We hypothesized that neural activity in this region would be required during the process of making effort-related choices on a trial-by-trial basis. To test this hypothesis, we trained mice as previously described and tested the effects of optogenetic silencing during discrete phases of our training and testing protocol (**Fig. 3D**). Following T-maze behavior, we tested the effects of our manipulation on overall mobility in an open field (OF) and in a more generalized test of effortful behavior using the tail suspension test (TST) (**Fig. 3D**). During T-maze testing, we made within-subject comparisons for light OFF vs. light ON trials that were pseudorandomly interleaved during each testing day (**Fig. 3F**).

**Figure 3.**
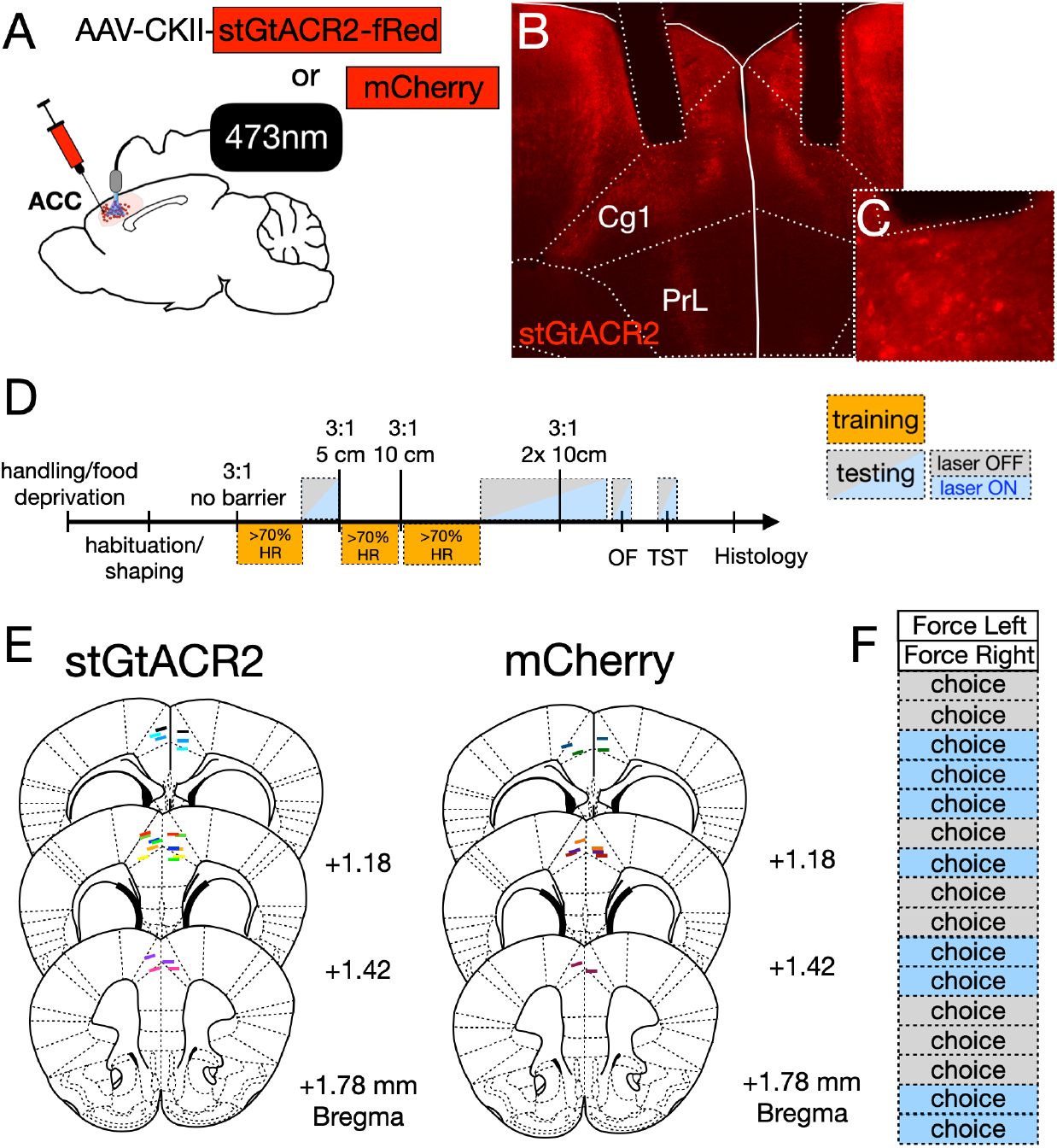
Experimental design and optic cannula for ACC silencing experiments. **A**) Bilateral ACC was injected with virus expressing an inhibitory opsin (stGtCAR2) or control (mCherry), and implanted with fiberoptic cannulae. **B**) Histology showing representative virus + cannula targeting (one cannula was implanted at a 15 degree angle). **C**) High magnification image showing membrane expression of stGtCAR2 under optic fiber. **D**) Overview of training and testing protocol for optogenetic silencing; optogenetic testing followed no barrier and/or barrier training phases, and during an open field (OF) and tail suspension test (TST). **E**) Targeting of optic cannulae. **F**) Example of a typical test day with 2 forced and 16 choice trials with light delivered in a pseudorandom pattern.

To test the acute necessity of ACC activity for EBD, we first examined the requirement for neural activity at discrete times during the trial cycle in our behavior. Mice were placed into the start box of the maze for 6 seconds before lifting the start gate to initiate a trial, with an ~60 second intertrial interval (ITI) (**Fig. 4A**). Light was delivered to silence the ACC during discrete phases of this cycle, and we compared the %HR choices for light OFF vs light ON trials. Light delivered during the trial itself was spatially restricted to cover the stem and choice point regions of the T-maze such that mice would need to commit to a choice while the ACC was silenced (**Fig. 4B**). We initially silenced the ACC from the end of the ITI through the choice point of the maze and observed a robust reduction in high effort, HR choices for light ON vs. OFF trials (**Fig. 4C**). We observed a similar reduction in HR choices when light was delivered during the ITI and in the start box, but not during the trial itself (**Fig. 4D**). However, there was no effect of silencing the ACC on %HR choices when light ON was limited to the home cage ITI (**Fig. 4E**). Notably, the largest effects were observed when ACC was silenced only during the choice part of the trial itself (between lifting the start gate and committing to a choice) (**Fig. 4F**).

**Figure 4.**
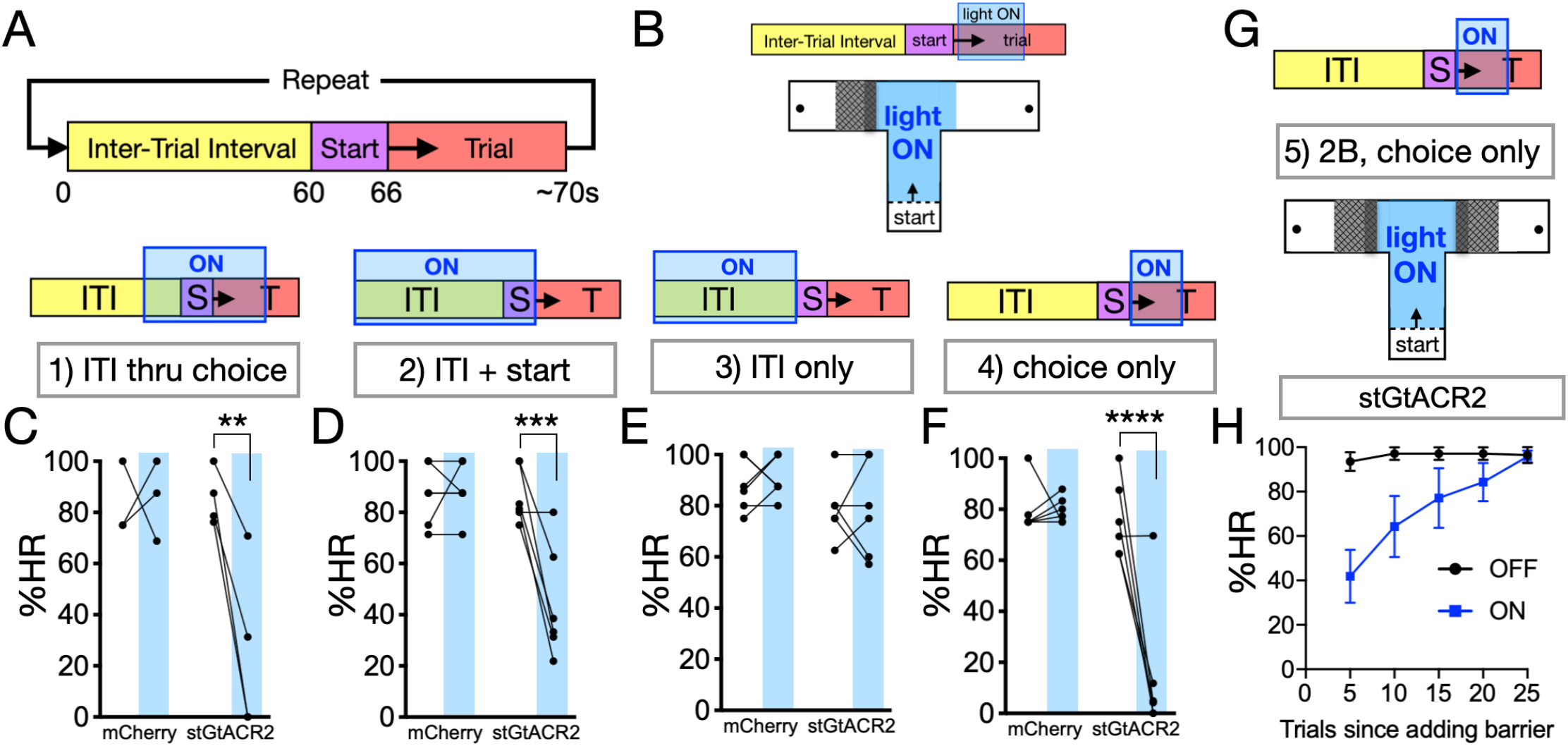
Optogenetic silencing of bilateral ACC reversibly disrupts high effort choices. **A**) Schematic representing trial cycle on which light OFF/ON epochs are overlaid. **B**) Example of light ON overlaid for choice only, light ON during trial itself extends spatially from start through choice point. **C**) Quantification of %HR choices for light OFF vs. ON, end of ITI through choice point, mCherry and stGtACR2; (blue shading is light ON) [*F_Interaction_*(1,10)=7.13, *P*=0.024, *F_Laser_*(1,10)=6.20, *P*=0.032, *F_Virus_*(1,10)=6.14, *P*=0.033], 2-way ANOVA with Sidak posthoc test. **D**) Quantification %HR for light OFF vs. ON, ITI + start only [*F_Interaction_*(1,20)=13.6, *P*=0.002, *F_Laser_*(1,20)=11.2, *P*=0.003, *F_Virus_*(1,20)=14.0, *P*=0.001], 2-way ANOVA with Sidak posthoc test. **E**) Quantification %HR for light OFF vs. ON, ITI only [*F_Interaction_*(1,20)=0.054, *P*=0.82, *F_Laser_*(1,20)=0.048, *P*=0.83, *F_Virus_*(1,20)=3.98, *P*=0.06], 2-way ANOVA with Sidak posthoc test. **F**) Quantification %HR for light OFF vs. ON, choice only [*F_Interaction_*(1,20)=20.8, *P*=0.0002, *F_Laser_*(1,20)=25.8, *P*=0.0001, *F_Virus_*(1,20)=20.6, *P*=0.0002], 2-way ANOVA with Sidak posthoc test. **G**) 2 Barrier control continuation for stGtACR2 choice only condition. **H**) %HR for 2 Barrier control, consecutive trials quantified in blocks of 5 for light OFF vs. ON. For (**C**), mCherry (n=3), stGtACR2 (n=4). For (**D-H**), mCherry (n=5), stGtACR2 (n=6); *p<0.05, **p<0.01, ***p<.001, ****p<0.0001.

There was no effect in mCherry-expressing, laser only mice for any of the conditions (**Fig. 4C-F**). These data demonstrate that the activity of ACC excitatory neurons is required on a trial-by-trial basis for mice to choose to exert greater effort for a larger reward, when a less effortful, lower reward alternative is available. Moreover, this activity is most critical during periods of time when mice are preparing for, evaluating and ultimately selecting an effortful action. The requirement for activity in the start box was somewhat surprising. Although we cannot rule out that this reflects some delay in re-establishing meaningful, task-related ACC representations after an extended period of silencing, this finding suggests that ACC activity may be important during a choice-planning phase immediately preceding the trial.

To better interpret these results, we equalized effort by adding a second barrier to the LR arm and continued to run trials with silencing during the trial itself (**Fig. 4G**). While mice initially persisted in choosing the LR arm when the ACC was silenced, they did adjust their behavior to select the more highly rewarded arm for both light OFF and ON trials (**Fig. 4H**). This suggests that absolute deficits in spatial recall, ability to enact an effortful response or motivation/preference for larger rewards cannot explain our results. The initial persistence in choosing the formerly low effort, LR arm is reminiscent of effects seen in ACC lesioned rats^7,28^. Because our manipulation is limited to the pre-choice period and light OFF trials are interspersed, it seems unlikely this reflects impaired updating of task models. However, ACC activity may be required to initially translate updated models into action selection.

To better understand the effects of silencing ACC on T-maze decision making we also conducted a separate experiment in which silencing was delivered in 3-day blocks during trials themselves (**Fig. 5A and B**). This design is more similar to how barrier T-maze experiments have typically been conducted, allowing better comparison with pharmacology or lesion studies. In this experiment, we initially silenced the ACC during three consecutive days in the no barrier training condition (**Fig. 5C**). There was minimal impact of silencing the ACC during choice trials that occurred before a barrier was introduced, although mice were slightly more likely to choose the LR arm when the light was ON (**Fig. 5C**). Like our other data, silencing ACC resulted in a large and highly significant disruption in high-effort choices after a barrier was introduced and this was stable over at least several days (**Fig. 5D**). When a second barrier was added to the maze, we again saw that silencing ACC continued to bias mice towards LR choices, but this effect was limited to the first day of the 2-barrier block (**Fig. 5E**). There was no effect of light alone in a similar 3-day block in mCherry expressing control mice indicating observed results are not due to effects of laser by itself (**Fig. 5F**). Taken together, these data support our conclusion that ACC activity is stably required during the evaluation and selection of actions which will incur an effort cost.

**Figure 5.**
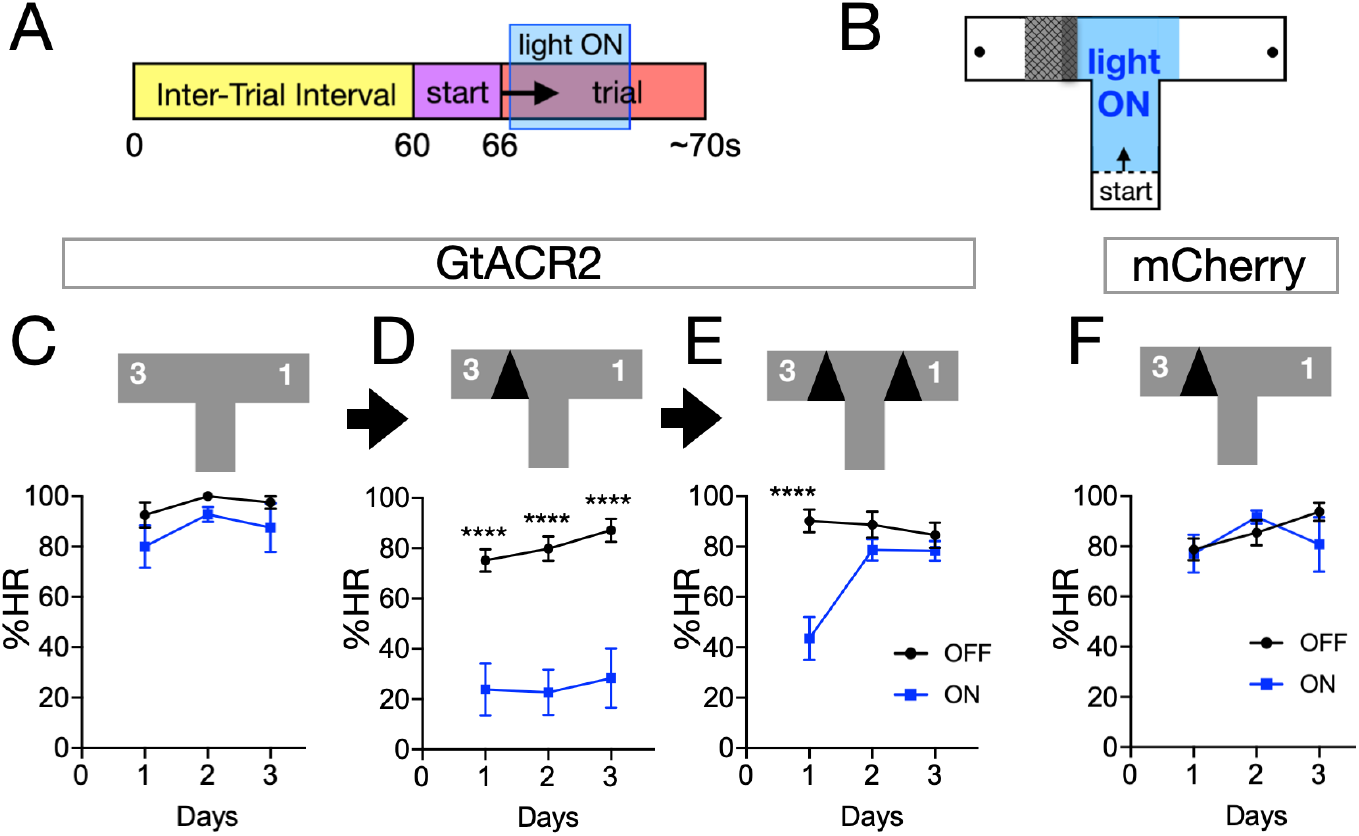
Optogenetic silencing of bilateral ACC reversibly disrupts high effort choices across days. **A-B**) Schematic for light ON vs. OFF during choice trials delivered across sequential 3 day blocks. **C**) Quantification of %HR choices for light OFF vs. ON, 3:1 differential no barrier condition [*F_Interaction_*(2,24)=0.10, *P*=0.904, *F_Days_*(2,24)=1.53, *P*=0.24, *F_Laser_*(1,24)=4.30, *P*=0.049], 2-way ANOVA with Sidak posthoc test (n=5). **D**) Quantification %HR for light OFF vs. ON, 3:1 one barrier condition [*F_Interaction_*(2,42)=0.11, *P*=0.89, *F_Days_*(2,42)=0.76, *P*=0.47, *F_Laser_*(1,42)=91.99, *P*<0.0001], 2-way ANOVA with Sidak posthoc test (n=8). **E**) Quantification %HR for light OFF vs. ON, 3:1 two barriers condition [*F_Interaction_*(3,38)=9.40, *P<0.0001, F_Days_*(3,38)=8.96, *P*=0.0001, *F_Laser_*(1,38)=11.34, *P*=0.002], 2-way ANOVA with Sidak posthoc test (n=7). **F**) Quantification %HR for light OFF vs. ON, mCherry control condition 3:1 one barrier condition [*F_Interaction_*(2,26)=1.20, *P=0.32, F_Days_*(2,26)=2.10, *P*=0.14, *F_Laser_*(1,26)=0.33, *P*=0.57], 2-way ANOVA with Sidak posthoc test (n=6). *p<0.05, **p<0.01, ***p<.001, ****p<0.0001.

### Silencing ACC disrupts choice trajectory during decision making

Our validation analyses revealed that choice trial duration is sensitive to changes in effort-reward tradeoffs (**Fig. 2**). Therefore, we predicted that a more detailed evaluation of choice behavior might reveal a more nuanced understanding of the role of ACC in EBD. A critical advantage of our assay, over operant tasks, is that task events are spatially and temporally segregated^45^. For example, evaluation of effort-reward tradeoffs and action selection occur prior to effort itself (climbing the barrier). A commonly employed alternative involves lever pressing in an operant chamber where mice must decide when to stop pressing a lever for a preferred reward over pursuing a less desired alternative. We hypothesized that the effects of silencing ACC activity on T-maze decision making would impact the time it takes mice to make choices in our maze and that these effects might be distinct for different zones within the maze. We used DeepLabCut^41^ to train a neural network for precise key point tracking of mice in our assay and custom software to analyze these data (**Fig. 6A**). Our lab recently developed BehaviorDEPOT, an open-source, flexible software package that allows customizable behavior analysis approaches based on machine-learning based tracking^42^.

**Figure 6.**
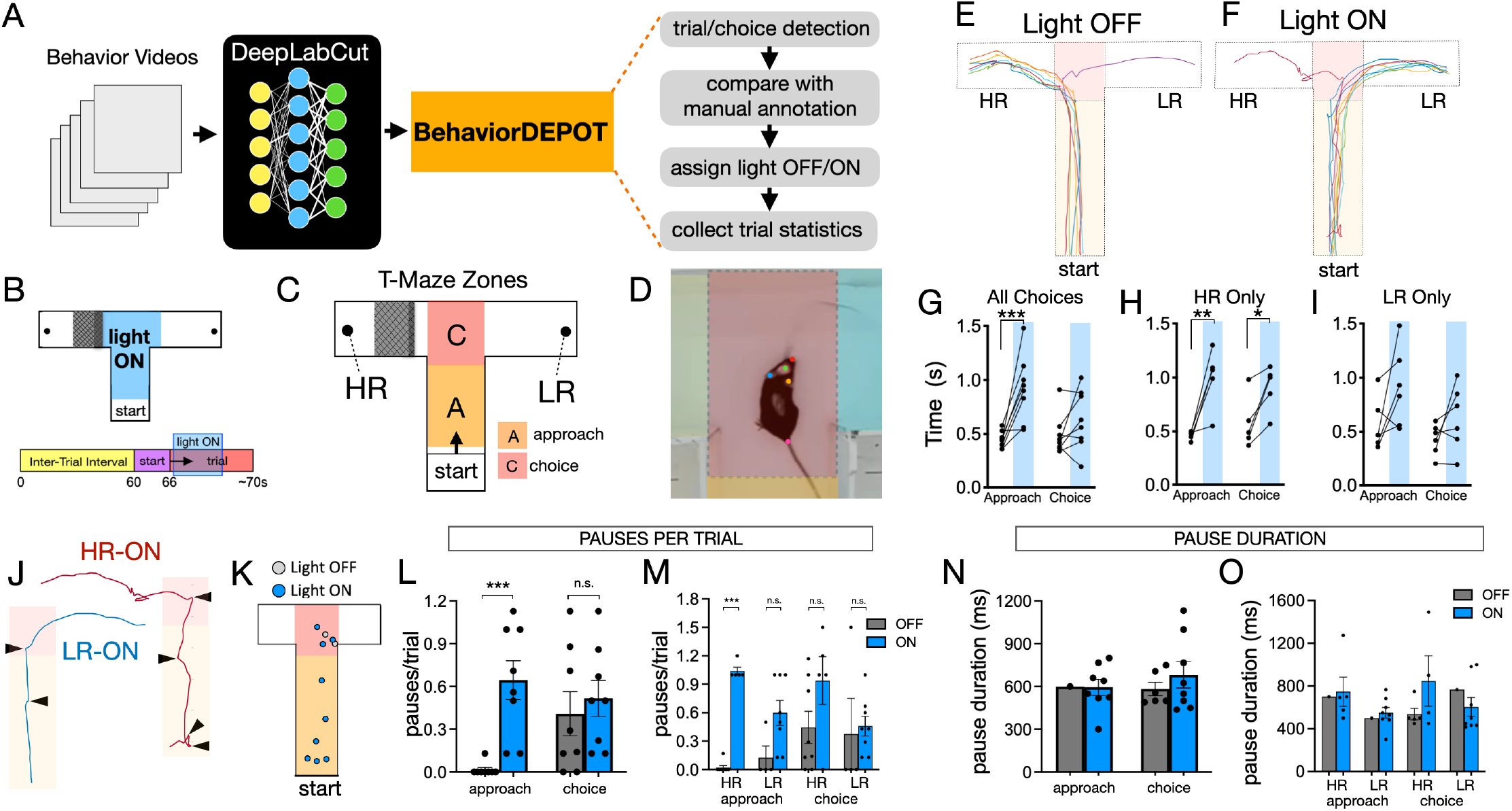
ACC silencing alters choice trajectories during effort-based action selection. **A**) Schematic representation of T-maze video processing: keypoint tracking with DeepLabCut (DLC) and analysis of tracking data using custom BehaviorDEPOT code. **B**) Behavior videos with silencing during action selection used for tracking analysis. **C**) Schematic representation of T-Maze zones assigned for subsequent analysis (‘approach’ + ‘choice’). **D**) DLC tracking for a mouse in the choice zone of the maze (head mounted implant used as tracking point (green dot)). **E**) Example trajectory plot for a mouse showing light OFF trials, and **F**) light ON trials with overlaid approach and choice zones. **G**) Quantification of time in approach and choice zones for light OFF vs. ON trials, one barrier condition - all choices [*F_Interaction_*(1,28)=4.41, *P*=0.045, *F_Zone_*(1,28)=2.47, *P*=0.13, *F_Laser_*(1,28)=14.35, *P*=0.0007], 2-way ANOVA with Sidak posthoc test. **H**) Quantification for HR choices only [*F_Interaction_*(1,16)=1.16, *P*=0.30, *F_Zone_*(1,16)=0.011, *P*=0.92, *F_Laser_*(1,16)=20.9, *P*=0.0003], 2-way ANOVA with Sidak posthoc test. **I**) Quantification for LR choices only [*F_Interaction_*(1,20)=0.47, *P*=0.50, *F_Zone_*(1,20)=3.58, *P*=0.07, *F_Laser_*(1,20)=6.05, *P*=0.02], 2-way ANOVA with Sidak posthoc test. **J**) Example, single trial trajectory plots for light ON condition (blue: low effort = ‘LR-ON’ and chrimson: high effort = ‘HR-ON’) with locations of micropauses marked (arrowhead). **K**) Locations of all pauses for example mouse. **L**) Quantification of pauses per trial for ALL choices, light OFF vs. ON [*F_Interaction_*(1,40)=8.30, *P*=0.006, *F_Laser_*(1,40)=11.35, *P*=0.0017, *F_Zone_*(1,40)=0.56, *P*=0.46], 2-way ANOVA with Sidak posthoc test, and **M**) broken down by choice type [*F_Laser_*(1,20)=35.12, *P<0.0001*, *F_Choice_*(3,40)=1.47, *P*=0.24, *F_LaserXChoice_*(3,20)=6.27, *P*=0.004], Mixed-effects analysis with Sidak posthoc test. **N**) Quantification of pause duration, ALL choices, light OFF vs. ON [*F_Interaction_*(1,27)=0.176, *P*=0.68, *F_Laser_*(1,27)=0.357, *P*=0.56, *F_Zone_*(1,27)=0.774, *P*=0.39], 2-way ANOVA with Sidak posthoc test, and **O**) broken down by choice type [*F_Laser_*(1,35)=0.803, *P=0.38*, *F_Choice_*(3,35)=0.318, *P*=0.81, *F_LaserXChoice_*(3,35)=0.168, *P*=0.92], Mixed-effects analysis with Sidak posthoc test. *p<0.05, **p<0.01, ***p<.001, ****p<0.0001.

We developed custom BehaviorDEPOT automated algorithms to process key point tracking data to segregate choice trials based on position as mice traverse zones sequentially during a choice. The specific sequence further defines whether the individual trials correspond to HR or LR choices, and we have validated performance of this process by comparison with manual annotations (**Fig. 6A**). We then assign the identity of trials as light OFF vs. light ON and automatically collect trial statistics segregated by zone and trial identity (light OFF/ON, HR/LR) (**Fig. 6A**). We processed and analyzed videos this way for testing days in which ACC was silenced during choice trials only (corresponding with ’approach’ and ’choice’ zones) (**Fig. 6B-D**). In this way, we can plot and analyze choice trajectories in the maze for light OFF vs. light ON conditions (**Fig. 6E-F**). We analyzed the time mice spent in the approach and choice zones of the maze segregated by light condition (OFF vs. ON) for all choices together and separated by choice type. When choice types were grouped, there was a highly significant increase in approach zone time for light ON vs. OFF trials, but not for the choice zone (**Fig. 6G**).

When we examined HR choices by themselves, time in approach and choice zones was significantly greater for the light ON condition, but we found no significant effects for LR choices (**Fig. 6H-I**). These data indicate that mice are slower to approach the maze choice point when ACC is silenced, particularly if they ultimately make a high effort choice. We next wondered what accounted for the increase time in the approach zone, which led us to examine choice trajectories in greater detail. We noticed that light ON trajectories were less smooth and that this was due to changes in the microstructure of ballistic movements (**Fig. 6J**). Specifically, mice exhibited brief micropauses that chunked movements into discrete segments when ACC was silenced (**Fig. 6K**). Video analyses revealed that micropauses were common in the approach zone when ACC was silenced, but almost never occurred during light OFF trials (**Fig. 6L**). In contrast, choice zone pauses were common for both conditions (**Fig. 6L**). When dividing by choice type, this difference was only significant for high effort choices (**Fig. 6M**). This indicates that approach zone micropauses occur on trials in which mice make a high effort choice despite silencing of ACC activity. There was no difference in micropause duration for any condition (**Fig. 6N-O**). Taken together, these results suggest that silencing ACC has the greatest effects on behavior in the approach zone, when mice may be evaluating or planning their choice. Thus, ACC may be required to support the efficiency or continuity of effortful action selection processes.

### Silencing ACC does not disrupt movement or effortful responses outside a decision-making context

Our results indicate that ACC activity is required for choosing to exert greater effort for a larger reward when a less effortful, lower reward alternative exists. We therefore wondered if silencing ACC would have more generalized effects on willingness or ability of mice to exert effort during behavior that is not explicitly reward-based or goal-directed. To investigate this, we examined the effects of silencing ACC during a tail suspension test (TST). The TST is a widely employed test for depression-related behavior in which mice are exposed to an inescapable acute stressor (tail suspension). Mice confronted with tail suspension stress either engage in bouts of effortful escape-related behavior or adopt an energy conserving immobile posture. While immobility has generally been taken as a measure of ’behavioral despair’ – a purportedly depression-like response – a more conservative interpretation is that both immobility and struggling can be considered potentially adaptive responses. In this case, we are not making any claims about relevance to depression and only using the assay as a measure of tendency to exert effort. Silencing neural activity in ACC during the TST did not reduce effortful struggling behavior (**Fig. 7A-B**). This suggests that silencing the ACC does not cause generalized reductions in effort outside the context of goal-directed decision making.

**Figure 7.**
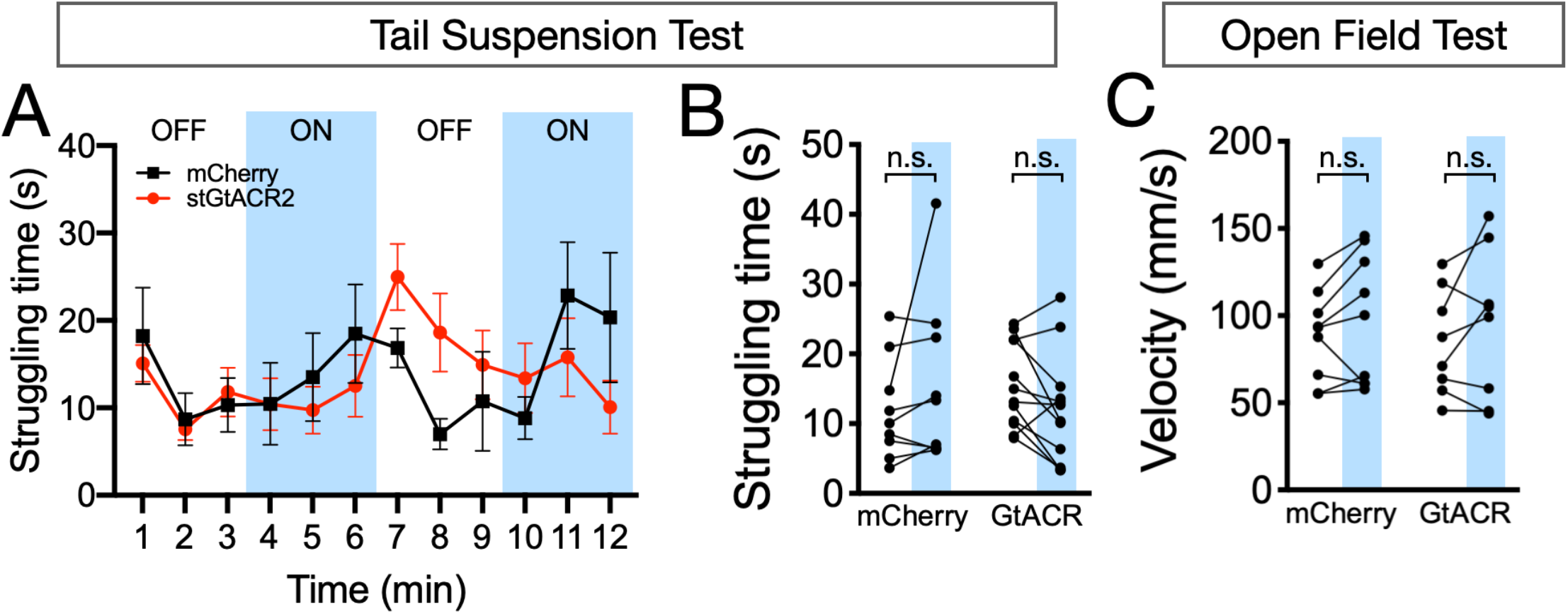
Optogenetic silencing of ACC has no effect on tail suspension test or open field tests. **A**) Quantification of mean struggling time calculated for each minute of the tail suspension test with optogenetic silencing of ACC (blue shading = light ON). **B**) Summary data for light OFF vs. ON for mCherry (n=9) and stGtACR2 (n=12) conditions [*F_Interaction_*(1,38)=1.96, *P*=0.17, *F_Virus_*(1,38)=0.0021, *P*=0.96, *F_Laser_*(1,38)=0.0028, *P*=0.96], 2-way ANOVA with Sidak posthoc test. **C**) Quantification of mean velocity for open field test with optogenetic silencing of the ACC, data for light OFF vs. ON for mCherry (n=9) and stGtACR2 (n=8) conditions [*F_Interaction_*(1,30)=0.0033, *P*=0.95, *F_Virus_*(1,30)=0.076, *P*=0.78, *F_Laser_*(1,30)=0.68, *P*=0.42], 2-way ANOVA with Sidak posthoc test. n.s. = not significant.

Our data also indicate that silencing ACC disrupts motoric aspects of behavior as mice approach a maze choice point. We therefore wondered whether this reflects generalized deficits in mobility, or is specific to situations that involve explicit decision making. To test this, we observed the effects of silencing the ACC when mice are exploring an open field (OF). We found that there is no difference in the average velocity of mice during light OFF vs. ON epochs for either stGtACR2 expressing mice or mCherry controls (**Fig. 7C**). This suggests that the observed disruptions in choice trajectory are specific to decision making, supporting the conclusion that they represent disturbances in goal-directed cognition.

## DISCUSSION

Here we report on the development and validation of a mouse version of the barrier T-maze assay. Mice rapidly modify their choices based on effort-reward tradeoffs and barrier climbing incurs a minimal time cost. Briefly silencing ACC excitatory neurons during action selection, or when mice are waiting to start a trial, rapidly and reversibly disrupts preference to exert greater effort for a larger reward. Instead, mice express a preference for a low effort, low reward alternative. This effect was absent when there was no effort cost, and reversed when effort cost was equalized, indicating that our results are not due to impairment in recall, capacity to perform the effortful action or preference for larger rewards. Silencing ACC also disrupts choice trajectory, causing mice to take longer to approach the maze choice point and select an action. This is at least in part due to changes in the microstructure of ballistic movements towards the choice point. Silencing ACC disrupts the typically continuous trajectory to the choice point, introducing frequent micropauses into the approach trajectory. In contrast, effortful behavior and overall movement were not impaired outside a goal-directed context. These findings indicate a role for ACC excitatory neurons in the evaluation and selection of effortful actions and online control of goal-directed action sequences.

The prefrontal cortex is postulated to exhibit a modular organization, but the role of ACC is extensively debated^46^. Prominent theories posit primary roles in resolving conflict, computing the value of effortful cognitive control over task performance and comparing the value of alternative actions in a decision oriented context^16,47,48^. Our data are consistent with prior lesion studies done in rats making effort-based choices^7,8,13,28^. However, we extend this work to define a role for ACC in computations that occur before and/or during effortful action selection. ACC neurons can dynamically encode potential actions with multiplexed representation of value, cost, and reward probability^49–52^. In rats, ACC activity in both single units and at the population level encode choice value that integrates reward magnitude and effort cost^11,14,15,36^. We find that silencing ACC during a choice, or when preparing for choice trials in the start box, disrupts effortful action selection. This may occur because ACC is required for the action-value associations which are necessary for making advantageous choices. Notably, this does not simply result in chance levels of performance, but instead biases mice towards low effort choices. This is consistent with computational models which propose that top-down ACC control is necessary to override effort-averse action selection systems in the striatum^53^.

In addition to changing choice preference, mice also exhibit changes in choice trajectory on trials where ACC was silenced. Beyond assigning effort-related value to actions, successful EBD requires monitoring and controlling action sequences required to reach the desired goal. Several studies find ACC population activity is involved in tracking or maintaining progress through a sequence of goal-directed actions^17,54,55^. If silencing ACC disrupts monitoring of progress towards a goal, then mice might need to pause and reorient frequently. The ACC is also intimately linked with motor output structures in the brain. ACC damage in patients can result in akinetic mutism, an absence of willed motor actions, and stimulation prompts reports of enhanced will to persevere through a challenge^56,57^. ACC neurons are particularly responsive to the organization or execution of goal-directed action sequences^58–62^. Thus, in addition to tracking progress, ACC may exert online control over the initiation and maintenance of goal-directed motor output responsible for lower level action sequences. Our results suggest a role for ACC in trial preparation, and as mice approach the choice zone and select an action. However, it is likely that ACC is important throughout a trial. One study found that ACC lesioned rats initially turned towards the barrier on the first few trials after a lesion, but failed to climb the barrier^29^. They suggest that ACC is important for maintaining effort while carrying an action sequence to completion. For the most part, we have not seen this type of behavior and mice typically turn directly to the low effort arm on trials where ACC was silenced. This is consistent with a role in processes occurring prior to action selection, though does not rule out important functions of ACC in carrying out effort or interpreting outcome.

Distinct cortical output pathways may segregate specific functions^19,63–65^. Our study expressed an inhibitory opsin driven by a promoter with some specificity for excitatory neurons (CaMKII), so our results may be driven by excitatory rather than inhibitory ACC neurons. A similar study using chemogenetics found that either activating or inhibiting ACC excitatory neurons disrupts effortful choices^32^. This suggests that increasing overall activity, if not timed precisely or coordinated appropriately amongst excitatory neurons, can also disrupt effortful choices. ACC projection neurons densely innervate dorsal striatum and mediodorsal thalamus^66^, but also target a number of other cortical and subcortical structures implicated in motivated behavior^67^. Lesions that disconnect ACC from basolateral amygdala or ventral striatum disrupt effortful choices in the barrier T-maze^9,68^ and other ACC targets including the habenula and anterior insula are also implicated in spatial EBD^69–71^. Moreover, dopaminergic input to both ventral striatum and ACC are important for EBD^26,27,72–76^. These data begin to define circuits interconnecting these regions that form distributed neural networks mediating EBD^45,77^. Future studies should investigate these cell-type specific circuits, by targeting genetically encoded opsins and/or calcium indicators. A deeper understanding of these ACC circuits will shed light on relevant neuropsychiatric disorders and their effective treatment.

## Acknowledgements

This work was funded by NIH K08MH116125 (SAW).

## Notes

**Conflict of interest**: The authors declare no competing financial interests

### Competing Interest Statement

The authors have declared no competing interest.

### Summary of Updates

Figure 6 has been revised to include new data about mouse trajectory during decision making. The text has been updated throughout to reflect these data and introduction/discussion have been revised. Author list has been updated.

